# Diffusion-weighted steady-state free precession imaging in the ex vivo macaque brain on a 10.5T human MRI scanner

**DOI:** 10.64898/2025.12.12.694017

**Authors:** Benjamin C. Tendler, Shaun Warrington, Mohamed K. Selim, Wenchuan Wu, Gregor Adriany, Edward J. Auerbach, Alexander Bratch, Hamza Farooq, Noam Harel, Sarah Heilbronner, Saad Jbabdi, Steve Jungst, Christophe Lenglet, Ana M.G. Manea, Steen Moeller, Franco Pestilli, Pramod K. Pisharady, Kamil Ugurbil, Matt Waks, Essa Yacoub, Stamatios N. Sotiropoulos, Karla L. Miller, Jan Zimmermann

## Abstract

Diffusion MRI provides a non-invasive probe of local fibre bundles and long-range anatomical connections to characterise the structural connectome. One way to achieve very high spatial resolution diffusion MRI data for connectivity investigations is to scan ex-vivo brains over many hours or days, ideally at ultra-high field strength to boost signal levels. However, conventional diffusion MRI acquisition techniques do not generally deliver good data quality for the challenging conditions of ex-vivo tissue, characterised by reduced diffusivities and relaxation times when compared to in vivo. In this work, we investigate the potential of the diffusion-weighted steady-state free precession (DW-SSFP) sequence for ex vivo diffusion imaging of the macaque brain using a 10.5 T human MRI scanner with a conventional (*G*_max_ = 70 mT/m) gradient set. SNR-efficiency optimisations incorporating experimental relaxation times demonstrate that the DW-SSFP sequence is predicted to achieve improved or similar SNR efficiency compared to a diffusion-weighted spin- and stimulated-echo sequence. Importantly, DW-SSFP can achieve this with the additional benefit of negligible geometric distortions, unlike conventional diffusion MRI using an echo-planar imaging readout. Using optimised DW-SSFP sequence parameters, we propose a protocol at 0.4 mm isotropic resolution using a two-shell multi-orientation protocol (effective b-values of 3200 s/mm^2^ and 5600 s/mm^2^). We fit the data using Tensor, Ball and 3-Sticks and Constrained Spherical Deconvolution signal representations. The results demonstrate high-quality diffusivity estimates across the entire brain with the ability to resolve multiple fibre populations in challenging crossing-fibre regions. The data will be made fully open source and multimodal as part of the Center for Mesoscale Connectomics, providing a resource for future connectivity investigations.

## Introduction

Diffusion MRI is the leading method to non-invasively measure structural connectivity and tissue microstructure in the human brain. Whilst the majority of connectomics investigations are conducted using in vivo diffusion MRI, there are two major reasons why ex vivo diffusion MRI is of interest. Firstly, by enabling scan times on the order of many hours or days, ex vivo diffusion MRI can provide datasets achieving higher spatial resolution to characterise detailed fibre architectures (Roebroeck et al., 2019; Schilling et al., 2025). Secondly, ex vivo diffusion MRI can serve as a bridge between in vivo diffusion MRI and microscopy techniques, sharing common signals with the former and tissue state with the latter. Datasets combining diffusion MRI with microscopy are powerful for investigating the origins of image contrast (Choe et al., 2012; Mollink et al., 2017) and support the development of improved tractography tools and other signal modelling approaches (Howard et al., 2019; S. Zhu et al., 2025).

Hardware advances for connectivity investigations have typically focused on developing MRI scanners with improved gradient performance (Huang et al., 2021; Van Essen et al., 2013). However, increases in static magnetic field strength offer a complementary theoretical boost in SNR (Le Ster et al., 2022; Vaughan et al., 2001), provided one can develop acquisition techniques that address reductions in *T*_2_, increased image distortions (arising from a more inhomogeneous *B*_0_) (Vu et al., 2015), and transmit (*B*_1_) inhomogeneity predicted at ultra-high field (de Moortele et al., 2005; Gallichan, 2018; Lagore et al., 2025). These challenges are exacerbated in fixed ex vivo tissue characterised by a reduced *T*_2_ and diffusion coefficient compared to in vivo (Shepherd et al., 2009), necessitating shorter diffusion-encodings and higher b-values to achieve equivalent SNR and diffusion contrast.

In recent years, diffusion-weighted steady-state free precession (DW-SSFP) (Figure 1a) (Kaiser et al., 1974; Le Bihan, 1988; Merboldt et al., 1989b, 1989a) has emerged as a powerful technique for ex vivo investigations at ultra-high field, achieving high SNR efficiency and strong diffusion weighting in fixed tissue (McNab et al., 2009; Miller et al., 2012). Previous work has demonstrated that the SNR-efficiency of DW-SSFP is predicted to increase with magnetic field strength (Foxley et al., 2014). Importantly, DW-SSFP uses a very short TR (∼20-40 ms) that is compatible with a heavily segmented readout acquiring a single phase encoding line (or a small number of phase encode lines) per TR. These readouts are associated with negligible image artefacts, in direct contrast to more conventional diffusion-weighted spin-echo (DW-SE) sequences (Figure 1b) that are only efficient with long readouts that corrupt image fidelity (e.g. EPI distortions or spiral blurring).

**Figure 1.**
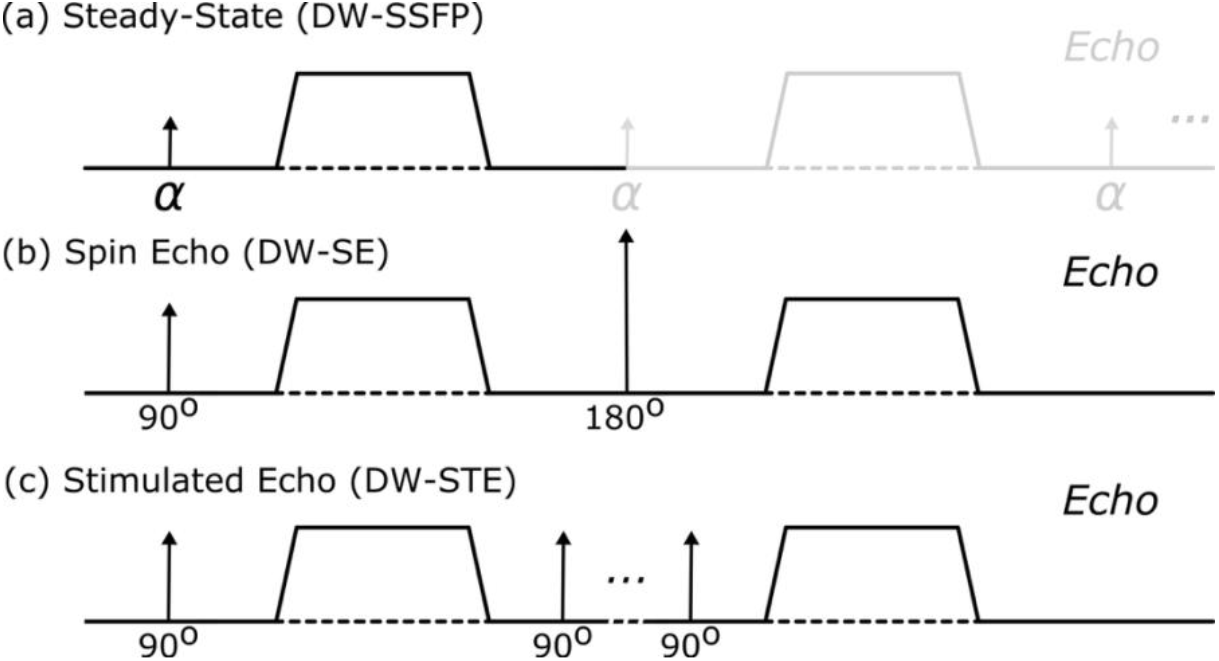
Diffusion encoding of different MRI sequences. (a) DW-SSFP, (b) DW-SE and (c) DW-STE. DW-SSFP (a) consists of a single RF pulse and diffusion gradient per TR (black line). A short TR and no spoiling of transverse magnetisation leads to magnetisation experiencing repeat sensitisation to RF pulses and pairs of dephasing/rephasing diffusion gradients over multiple TRs (grey line), consistent with more conventional diffusion MRI sequences (b and c). DW-STE (c) achieves strong diffusion-weighting with reduced *T*_2_ signal loss by increasing diffusion sensitisation longitudinally (associated with slow *T*_1_ recovery), beneficial for ultra-high-field imaging (a short *T*_2_ and long *T*_1_ imaging environment). This results in increased experimental time and signal-forming mechanisms that lead to a 2-fold reduction in signal levels when compared to a DW-SE.

Whilst DW-SSFP’s severe motion sensitivity limits in vivo investigations, this is not a confound for ex vivo acquisitions. When considering tractography investigations, the sequence’s complicated signal forming mechanisms have been addressed with advances in DW-SSFP signal modelling (McNab et al., 2009; Miller et al., 2012; Tendler et al., 2020). Whilst accurate diffusion modelling requires information about voxelwise tissue relaxation times and transmit inhomogeneity, this can be addressed by incorporating *T*_1_, *T*_2_, and *B*_1_ mapping techniques into data acquisition.

When considering ex vivo investigations, previous work with DW-SSFP has typically focused on imaging large, immersion-fixed samples including whole human (McNab et al., 2009; Tendler et al., 2022), primate (Bryant et al., 2021; Roumazeilles et al., 2020), and cetacean (Berns et al., 2015) brains, alongside human brain hemispheres (Cardenas et al., 2017; Edlow et al., 2018; Nolan et al., 2021; van Veluw et al., 2019; Wilkinson et al., 2016; Q. Zhu et al., 2025) on 3 or 7 T human systems. These acquisitions have demonstrated the potential of DW-SSFP for ex vivo investigations, routinely achieving sub-mm resolution with high-levels of diffusion contrast, even when limited to human MRI scanners with conventional gradient sets. Resulting datasets can be integrated into tractography pipelines, with improved SNR-efficiency and tract reconstructions at 3T in comparison to the DW-SE (Miller et al., 2012), and at 7 T in comparison to 3 T (Foxley et al., 2014) motivating the use of ultra-high field systems for DW-SSFP driven connectivity investigations.

In this manuscript, we investigate the potential of DW-SSFP on a bespoke 10.5 T human MRI scanner with a conventional (70 mT/m) gradient set for imaging a whole, perfusion-fixed ex vivo macaque brain. The scanner is equipped with a custom 40-channel receive coil designed for ex vivo macaque brains, providing improved brain coverage in comparison to conventional preclinical systems and minimal coil- to-tissue distance to boost SNR. Imaging a smaller perfusion-fixed sample is relatively unexplored in DW-SSFP, offering improved tissue preservation with anticipated reduced sensitivity to *B*_1_ inhomogeneity in comparison to existing high-field DW-SSFP studies.

We demonstrate that for ex vivo imaging at 10.5 T, DW-SSFP offers low-distortion imaging with high SNR efficiency over a wide target b-value range, predicting higher or similar SNR efficiency in comparison to a DW-SE and diffusion-weighted stimulated-echo (DW-STE) (Figure 1c) sequence. The proposed DW-SSFP acquisition and image reconstruction scheme achieves high SNR diffusivity estimates in an ex vivo macaque brain, with the acquired 400 μm (isotropic) dataset resolving multiple fibre bundles in complicated crossing-fibre regions.

This work is undertaken as part of the Center for Mesoscale Connectomics (https://mesoscale-connectivity.org), a collaborative effort acquiring a unique dataset combining in vivo MRI, ex vivo MRI and polarisation sensitive optical coherence tomography (PS-OCT) in whole macaque and human brains. The macaque imaging component benefits from in vivo and ex vivo data being acquired in the same brain sample, alongside complementary tract tracing and whole brain light sheet microscopy data, facilitating cross-domain, cross-scale comparisons. The methods development presented here is anticipated to form the blueprint for future ex vivo human brain acquisitions, where DW-SSFP has previously demonstrated considerable advantages, and the size of the human brain necessitates the use of a human MRI scanner as opposed to using preclinical systems with considerably higher gradient strengths.

### Theory

#### Diffusion-weighted steady-state free precession (DW-SSFP)

The diffusion encoding module of the DW-SSFP sequence consists of a single RF pulse (flip angle *α*, typically ≪ 90°) followed by a single diffusion-weighting gradient (length scale *q*^−1^), after which signal is acquired (Figure 1a – black line). The sequence is characterised by a short TR (relative to transverse relaxation times) and no spoiling of transverse magnetisation at the end of each TR. Magnetisation components persist over multiple TRs and experience repeat exposure to RF pulses and pairs of dephasing/rephasing diffusion gradients (Figure 1a – grey line) consistent with more conventional diffusion MRI sequences (Figures 1b and c), leading to a diffusion-weighted signal.

The signal-forming mechanisms of DW-SSFP lead to an inherently more complicated signal expression (Appendix 1) (Freed et al., 2001) in comparison to the DW-SE and DW-STE sequence (Appendix 2 and 3). When considering diffusion systems consisting of multiple Gaussian compartments, the DW-SSFP signal can be accurately described as the weighted-sum of magnetisation components associated with different b-values (Buxton, 1993; McNab & Miller, 2008; Tendler, 2025). From the perspective of extended phase graphs (EPG), this reflects the sum of individual magnetisation pathways associated with different evolution histories (Weigel, 2015). For a review of the DW-SSFP sequence, we recommend the article by (McNab & Miller, 2010). For a detailed description of the signal-forming mechanisms of DW-SSFP, we recommend the article by (Tendler, 2025).

The signal-forming mechanisms of DW-SSFP offer several advantageous properties for diffusion MRI investigations. Firstly a large fraction of each TR can be dedicated to the sequence’s readout, leading to high-SNR efficiency (Miller et al., 2012). Secondly, excited magnetisation components experience repeat sensitisation to diffusion gradients and are associated with long effective diffusion times (dictated predominantly by the *T*_1_ coefficient), achieving strong diffusion-weighting even when limited to conventional gradient sets. Thirdly, the short TR of the sequence means that echo times remain low, boosting signal levels in short *T*_2_ tissues. Fourthly, the short TR of the sequence means a single-line readout can be used whilst keeping acquisition times manageable, leading to datasets with minimal geometric distortions.

#### SNR Efficiency

SNR efficiency describes the amount of signal available from a sequence normalised by the effect of scan duration (Miller et al., 2012), defined as:

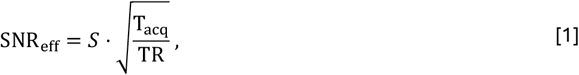

where SNR_eff_ = SNR efficiency, *S* = signal amplitude and T_acq_ = readout duration. SNR efficiency measures facilitate comparisons of the relative SNR available with different acquisition parameters and imaging sequences, with higher SNR_eff_ corresponding to a higher SNR per unit time.

The expression of *S* for the DW-SSFP sequence, *S*_SSFP_, is defined in Appendix 1, with the signal amplitude dependent on several sequence parameters (*G* = diffusion gradient amplitude, *δ* = diffusion gradient duration, *α* = flip angle, TR/TE = repetition/echo time), relaxation times 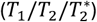 and the diffusion coefficient (*D*).

For the DW-SSFP sequence T_acq_ is defined as:

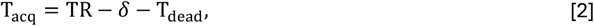

where T_dead_ defines the minimum time between the diffusion gradient and readout, fixed to 5 ms here for all simulations. Equivalent expressions for the DW-SE and DW-STE sequence are provided in Appendices 2 and 3.

## Methods

### Ethics

Experimental procedures were carried out in accordance with the University of Minnesota Institutional Animal Care and Use Committee and the National Institute of Health standards for the care and use of non-human primates (NHPs). All subjects were fed ad libitum within a light and temperature-controlled colony room. Animals had access to ad lib water.

### Brain sample and preparation

Two months post AAV injection for axonal and neuronal viral labelling, animals are euthanized using pentobarbital following ketamine/dexmedetomidine sedation and transcardially perfused using PBS followed by 4% hydrogel monomer solution (40% w/v acrylamide, 2% w/v bisacrylamide, 10X PBS, 8% w/v paraformaldehyde and distilled water). Brains are extracted and kept in 4% HMS for one week, and subsequently transferred to PBS and stored in a refrigerator until imaged (Shepherd et al., 2009).

Prior to imaging, brains are brought to room temperature and transferred into a form fitting shell (Figure 2) filled with a susceptibility matched electronic liquid (3M Flourinert FC-3283). Small air bubbles are removed using rotational tilting as well as using a soft catheter tip. Brains are left to sit overnight to alleviate any previously undetected air bubbles and then transferred to the magnet room on the morning of the scan. After ex-vivo MR data acquisition, specimens are transferred back into PBS and stored in the refrigerator. Following data quality controls specimens are transferred to the PS-OCT part of the project and sectioned into 300 micron thick slices.

**Figure 2.**
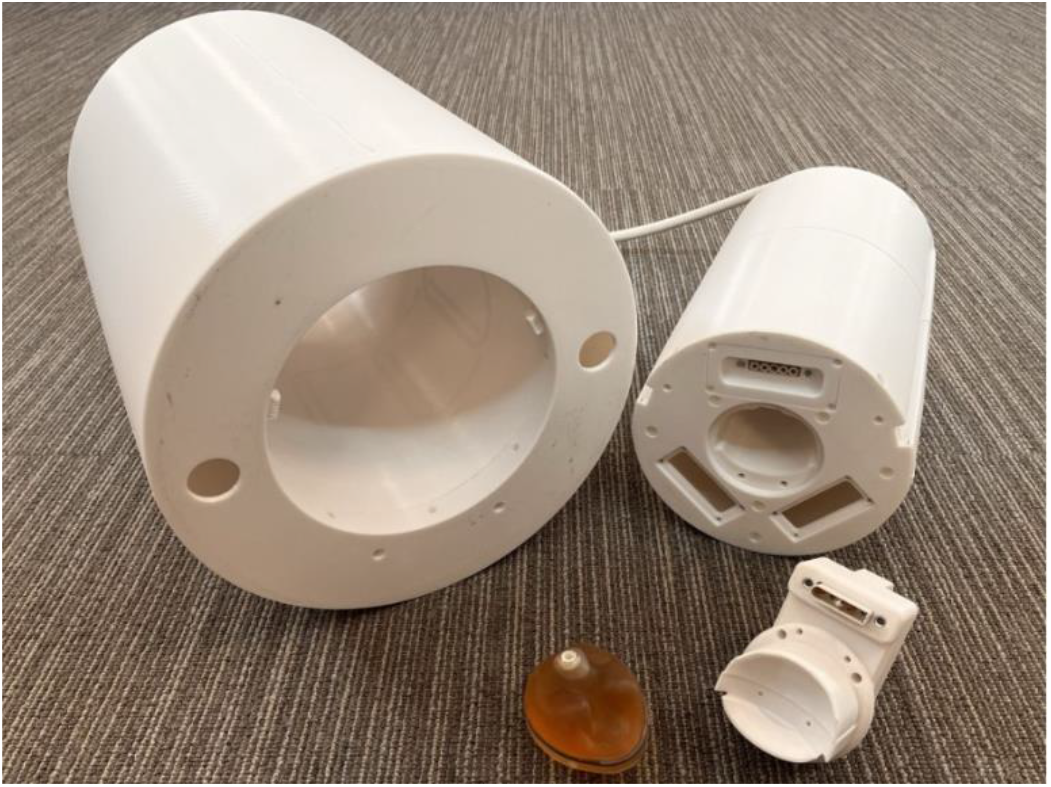
Overview of the RF Coil. The RF coil was custom designed for ex vivo macaque brain imaging, consisting of 8 transmit and 40 receive elements. The location of coil elements was chosen to provide full spatial coverage with minimal coil-to-tissue distance over the entire macaque brain, providing a considerable advantage in comparison to more conventional human and preclinical imaging coils that necessitate anatomical constraints. The macaque brain was secured inside a custom 3D printed shell to minimise sample motion and non-linear tissue deformities (e.g. shearing effects) over the time course of the acquisition. For a more detailed overview of the coil properties, see [Waks et al., 2025].

### MRI scanner and hardware

1x ex vivo macaque brain was scanned on Siemens MAGNETOM 10.5 T scanner (software version VE12U) based at the University of Minnesota (Center for Magnetic Resonance Research). The scanner was equipped with a SC72D gradient set (*G*_Max_ = 70 mT/m, G_Slew_ = 200 T/m/s). Data were acquired using a multi-channel (8 Tx, 40 Rx) coil custom designed for ex vivo macaque brain imaging (Figure 2). The coil is capable of 8 channel pTx but was used in conjunction with a 8-way splitter (Werlatone, Brewster, NY) in ‘single channel -transmit’ mode (i.e. each transmit coil element was presented an identical RF pulse with 45 deg phase shift between neighbouring channels).

### Sequence optimisation

Sequence parameters for the DW-SSFP acquisition were obtained using a custom optimisation routine designed to maximise SNR efficiency (see Theory) for a target b-value. As the DW-SSFP sequence does not have a well-defined b-value, an ‘effective’ b-value was used based on the level of predicted signal attenuation, defined as (Miller et al., 2012):

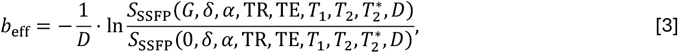

where the numerator and denominator of Eq. [3] are equivalent to expressions of the diffusion and non-diffusion weighted DW-SSFP signal (Appendix 1).

Parameter optimisation was performed in MATLAB (version 2023a) using the *lsqnonlin* algorithm, identifying a target *G, δ, α*, and TR that maximised SNR efficiency for a target b-value, *b*_target_, via:

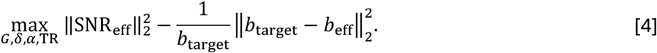

The optimisation incorporated experimental relaxation times (*T*_1_, *T*_2_) estimates from the ex vivo brain sample, alongside approximating the 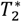 (50% of the *T*_2_) and a diffusion-coefficient (*D* = 2 · 10^−4^ mm^2^/s) to reflect ex vivo conditions in the target b-value range (1,000 to 10,000 s/mm^2^). To avoid gradient duty cycle limitations, we defined the maximum diffusion gradient amplitude, *G*_max_ = 52 mT/m based on previous optimisation work using an equivalent gradient set (Tendler et al., 2020). A maximum readout duration was fixed to 30 ms. Following the optimisation, a small correction was performed *δ* to ensure the applied diffusion gradient led to 2*π* · *n* dephasing at the target sample resolution (0.4 mm isotropic).

The relative SNR efficiency of the DW-SSFP sequence was compared with the DW-SE and DW-STE sequence (expressions for *S* and T_acq_ provided in Appendices 2 and 3) using an equivalent optimisation routine and limits (*G*_max_ = 52 mT/m, maximum readout duration = 30 ms).

### Acquisition protocol

Pilot experimental relaxation estimates in the ex vivo brain at 10.5 T (based on an inversion recovery and a multi-echo spin echo sequence) estimated an average *T*_1_ = 1350 ms and *T*_2_ = 28 ms. Based on the SNR-efficiency optimisations using these relaxation times, a two-shell acquisition was performed with *b*_eff_ = 3200 and 5600 s/mm^2^, alongside b_0_ volumes at a resolution of 0.4 mm isotropic. The highest b-value shell represents a trade-off between achieving sufficient acquisition SNR and diffusion contrast. The lower b-value shell was achieved by reducing the diffusion gradient amplitude (fixing all other parameters), ensuring the q-value achieved 2*π* · *n* dephasing at the target sample resolution (0.4 mm isotropic). Sequence parameters are provided in Table 1.

**Table 1:**
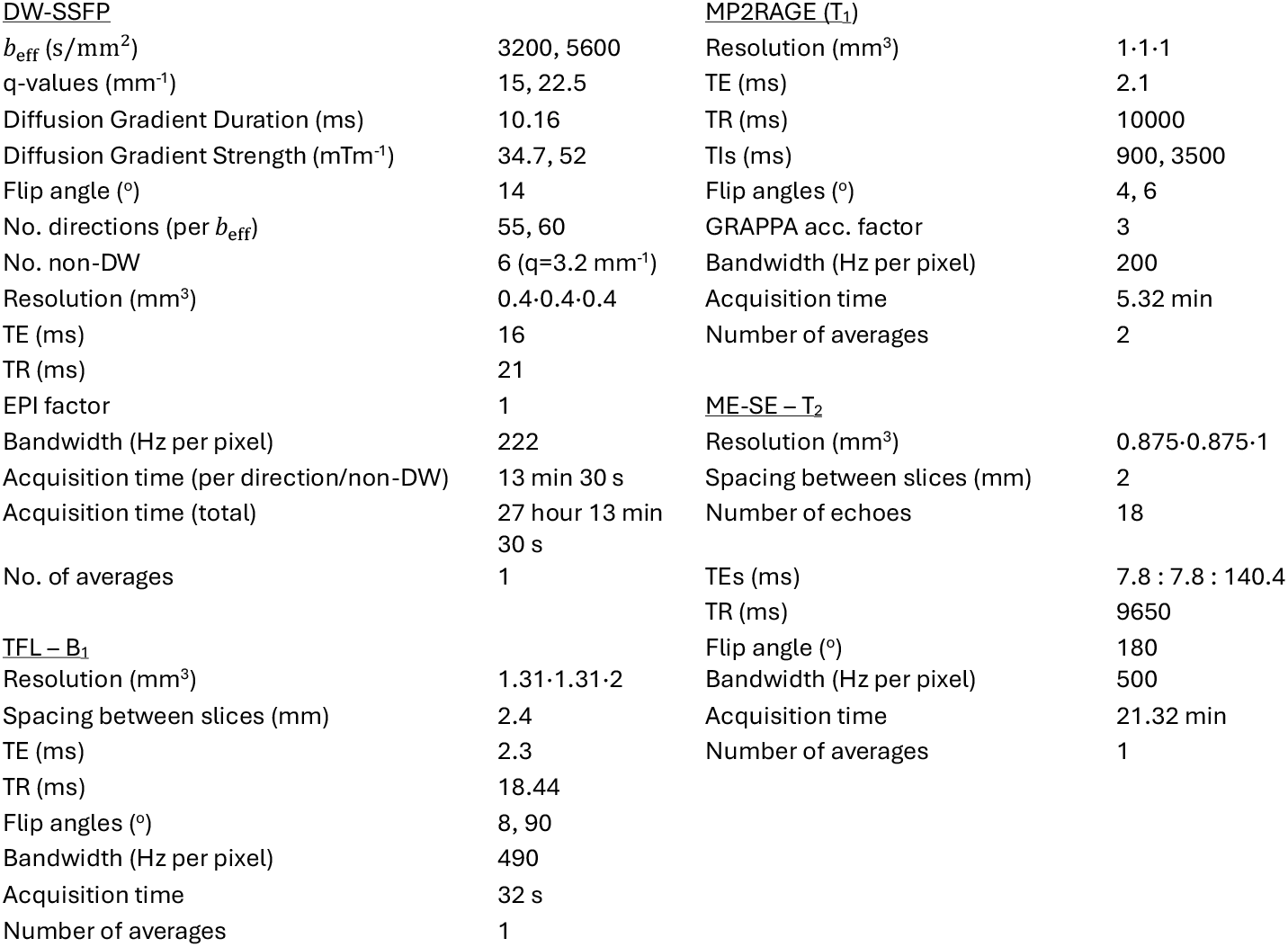
MRI acquisition parameters. The imaging parameters of the DW-SSFP sequence were based on the SNR-efficiency optimisation (see Methods). A quantitative *T*_1_ map was reconstructed from the Magnetization Prepared 2 Rapid Acquisition Gradient Echoes (MP2RAGE) data based on the scanner’s online reconstruction. The *B*_1_ map was reconstructed from the TFL data based on the scanner’s online reconstruction A quantitative *T*_2_ map was reconstructed from the ME-SE data using a custom extended-phase graphs (EPG) fitting routine similar to the approach in (Tendler et al., 2021).

The b_0_ volumes were interspaced throughout the acquisition to estimate and correct for signal drift over the time course of the acquisition, often observed in lengthy ex vivo acquisition protocols and typically attributed to gradient heating. For the b_0_ acquisitions, a small spoiler gradient is required to prevent the formation of banding artefacts associated with balanced SSFP sequences. Apart from this gradient, the b_0_ volumes were acquired with matched sequence parameters to the diffusion weighted volumes, setting *G* = 7.4 mT/m (*q* = 3.2 mm^−1^), leading to an effective b-value of 200 s/mm^2^. The q-value of 3.2 mm^−1^ was slightly higher than the q-value required to achieve 2*π* dephasing (2.5 mm^−1^), determined heuristically on the scanner to ensure the removal of all banding artefacts.

Accurate diffusivity estimation in DW-SSFP requires estimation of voxelwise *T*_1_ and *T*_2_ tissue relaxation times, alongside *B*_1_ transmit estimates. This can be appreciated in the signal model given in Appendix 1, where the relaxation and flip angle terms are not separable from the diffusion-weighted terms. Quantitative *T*_1_, *T*_2_ and *B*_1_ maps were obtained via a Magnetization Prepared 2 Rapid Acquisition Gradient Echoes (MP2RAGE), a multi-echo spin-echo (ME-SE) and turbo flash (TFL) sequence, with parameters and a description of the quantitative mapping approaches provided in Table 1. Quantitative maps were aligned to the space of the DW-SSFP data using a rigid body (6 degrees of freedom) coregistration with FSL FLIRT (Jenkinson et al., 2002; Jenkinson & Smith, 2001).

### Data processing and modelling

The DW-SSFP data were reconstructed offline (SENSE1) (Sotiropoulos et al., 2013), denoised in the complex domain (NORDIC) (Moeller et al., 2021) and corrected for signal drift over the time course of the acquisition. A rigid body (6 degrees of freedom) coregistration was performed between volumes using FSL FLIRT. Quality control (QC) measures were established using the EDDY QC toolbox, where we report the SNR measures from the b_0_ volumes and angular contrast to noise (CNR) for the diffusion-weighted volumes (Bastiani et al., 2019).

Signal modelling was performed incorporating a (1) Tensor, (2) Ball & 3-Sticks and (3) Constrained Spherical Deconvolution (CSD) signal representations. The Buxton DW-SSFP signal model (Buxton, 1993) was incorporated into a Tensor and Ball and 3-Sticks signal representation, with fitting performed using Markov Chain Monte Carlo (MCMC) with cuDIMOT (Hernandez-Fernandez et al., 2019), a GPU-driven signal modelling software package. Fifty posterior samples were taken per estimated parameter, with the Tensor estimated solely from the *b*_eff_ = 3200 shell, and Ball & 3-Sticks estimated from both shells assuming a single diffusion coefficient. Example tractography outputs were estimated from the Ball and 3 Sticks parameter outputs using FSL-XTRACT (Assimopoulos et al., 2024; Warrington et al., 2019).

CSD was performed using MRtrix3 (Tournier et al., 2019). Briefly following conversion of the pre-processed DW-SSFP data into an MRtrix-compatible format, multi-shell, multi-tissue CSD was applied. Tissue-specific response functions for white matter, grey matter, and cerebrospinal fluid were estimated in an unsupervised manner using the approach described in (Dhollander et al., 2016). Fibre orientation distributions (FODs) were then computed for each voxel using the three-tissue response functions.

## Results

Figure 3 compares the relative SNR efficiency predicted for the DW-SSFP, DW-SE and DW-STE sequence. The DW-SSFP sequence demonstrates higher SNR efficiency across the investigated b-value regime based on the pilot relaxation values. Importantly, while this reflects the quality of the data per voxel, the single-line readout of DW-SSFP offers the additional benefit of minimal image distortions even in the presence of field inhomogeneities, which can be a considerable challenge for conventional EPI-based DW-SE methods at ultra-high field. Supporting information Table S1 provides the estimated SNR-efficient sequence parameters for each sequence across the b-value range.

**Figure 3.**
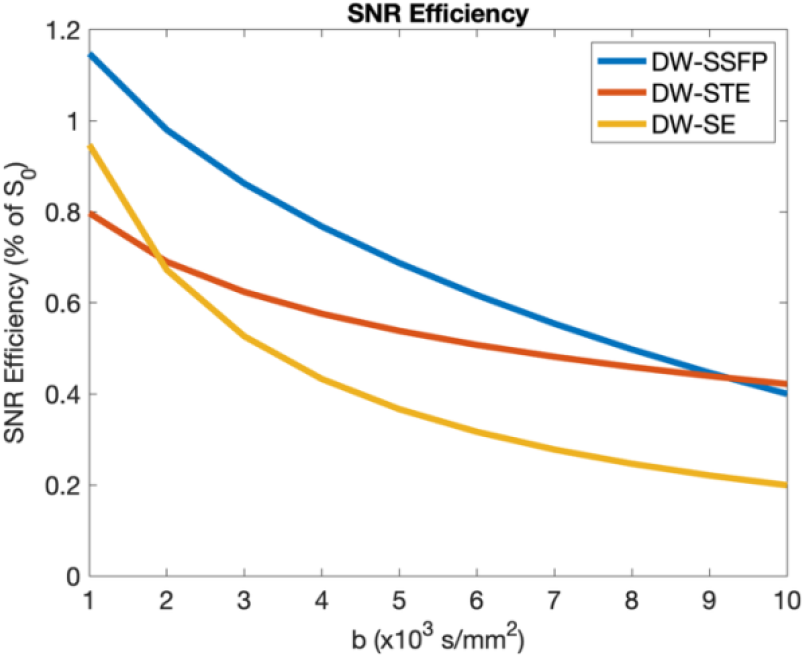
SNR efficiency comparisons. Optimised SNR-efficiency estimates for the DW-SE (yellow), DW-STE (red) and DW-SSFP (blue) sequence. DW-SSFP predicts increased or similar SNR efficiency across the investigated regime, closer to the DW-SE sequence at low b-values and the DW-STE sequence at high b-values. Optimisations based on scanner properties and pilot relaxometry estimates (*T*_1_ = 1350 ms and 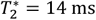) in the fixed ex vivo brain (see Methods). To aid visualisation the figure does not incorporate the impact of diffusion-weighting on SNR efficiency estimates, which is equivalent for all three sequences. No 2*π* · *n* dephasing correction was performed for the DW-SSFP SNR efficiency optimisation presented here.

Figure 4 displays example diffusion-weighted volumes acquired at *b*_eff_ = 5600 s/mm^2^ with orthogonal diffusion gradient orientations. Connectivity investigations are driven by differences in orientation-dependent contrast, and here we can visualise distinct anatomical contrast across the three orientations (example differences highlighted with the purple arrows), where we note that no geometric distortion correction has been applied to these data.

**Figure 4.**
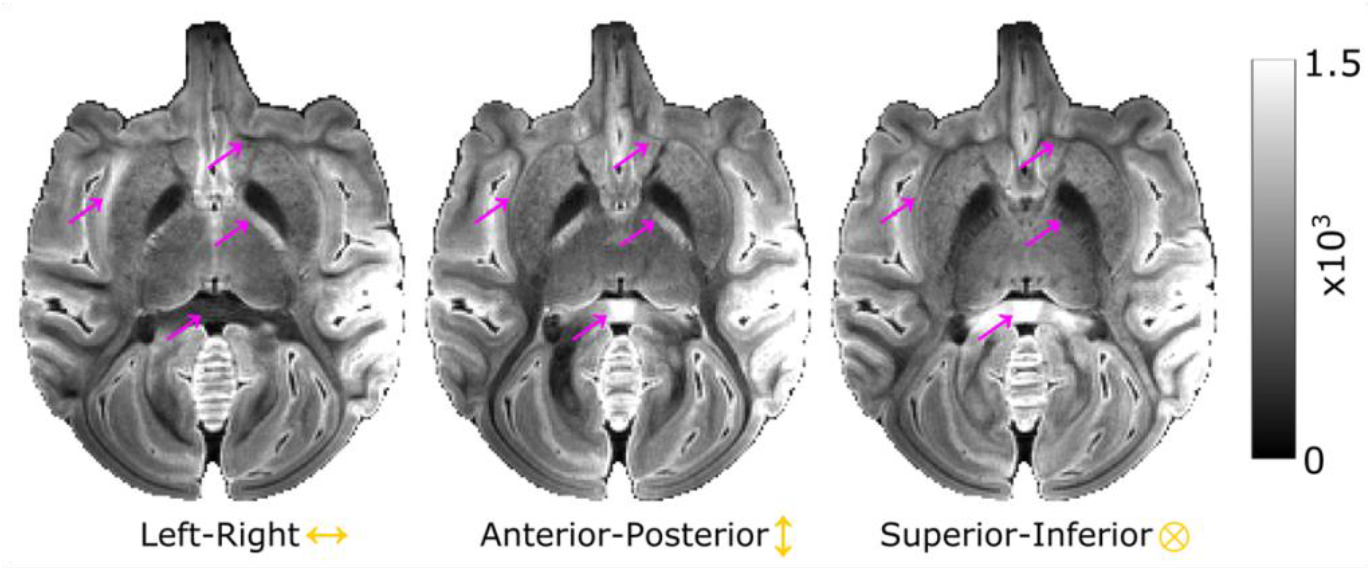
Orientation dependent diffusion contrast. Here we display three diffusion-weighted volumes acquired with *b*_eff_ = 5600 s/mm^2^ with the diffusion gradients oriented principally along the left-right (left), anterior posterior (middle) and superior-inferior (right) brain axis. Distinct anatomical contrast is present across the three images, with example differences highlighted with the purple arrows. Specific diffusion gradient orientations: Left-right = [0.975, 0.150, 0.165], anterior-posterior = [-0.005, 0.996, -0.087], superior-inferior = [-0.093, 0.218, 0.972].

Figure 5 displays the b_0_ and diffusion-weighted volumes (top row) averaged across all repeats/diffusion directions, alongside the CNR maps (bottom row) averaged across all diffusion directions based on the QC analysis. The diffusion-weighted volumes display excellent contrast, with the expected increase in diffusion-weighting as a function of effective b-value. Following offline reconstruction and denoising, we estimate an average SNR in the b_0_ volumes of 34.91, and CNR of 1.41 (*b*_eff_ = 3200 s/mm^2^) and 1.77 (*b*_eff_ = 5600 s/mm^2^) respectively. The average diffusion attenuation across the brain was 0.4 ± 0.1 (*b*_eff_ = 3200 s/mm^2^) and 0.24 ± 0.09 (*b*_eff_ = 5600 s/mm^2^). No correction for geometric distortion has been performed.

**Figure 5.**
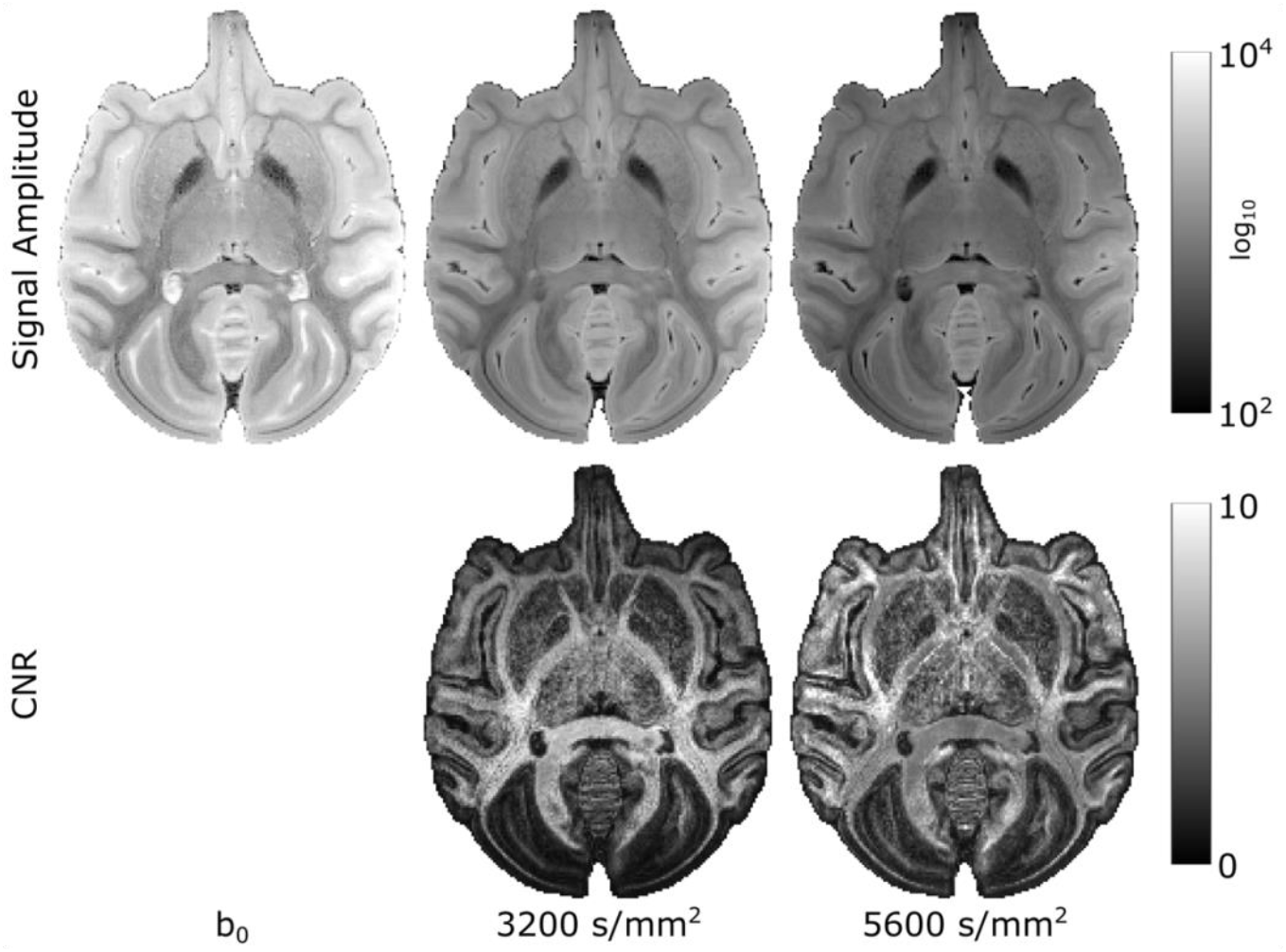
SNR and CNR. The top row displays the relative signal amplitude (log_10_ scale) for the b_0_ and diffusion-weighted volumes, here averaged across all repeats (b_0_) or diffusion directions (diffusion-weighted volumes) per b-value. Detailed spatial contrast is present across the brain even at the highest b-value. The bottom row displays the voxelwise CNR estimates averaged across all diffusion directions derived from EDDY QC, displaying high CNR across the brain, most predominantly in major white matter tracts.

The median relaxation times over the whole brain at the time of acquisition were estimated as *T*_1_ = 1050 ms, *T*_2_ = 53 ms and 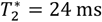. Note that these values are considerably different to the initial relaxation estimates used for the sequence optimisation (*T*_1_ = 1350 ms, *T*_2_ = 28 ms and 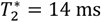), a likely result of the increased soaking time in PBS at the time of dataset acquisition. We discuss the consequences of this in terms of SNR-efficiency further in the Discussion.

SNR-efficiency and diffusion-weighting are strongly dependent on the flip angle in DW-SSFP, with reduced flip angles (relative to the CNR optimal flip angle) typically corresponding to a sharp drop in CNR (Figure 6a – blue line). Substantial *B*_1_ inhomogeneity at ultra-high field can lead to large spatial variations in SNR and diffusion-weighting across the brain, previously requiring the use of specialised multi-flip angle acquisition schemes (Tendler et al., 2020) or parallel transmit systems (Fritz et al., 2016) to address. The *B*_1_ inhomogeneity was found to be moderate across the macaque brain (Figure 6), a reflection of its size (maximum macaque brain dimension ∼ 7.5 cm) relative to the RF wavelength (∼ 7 cm in vivo) at 10.5 T.

**Figure 6.**
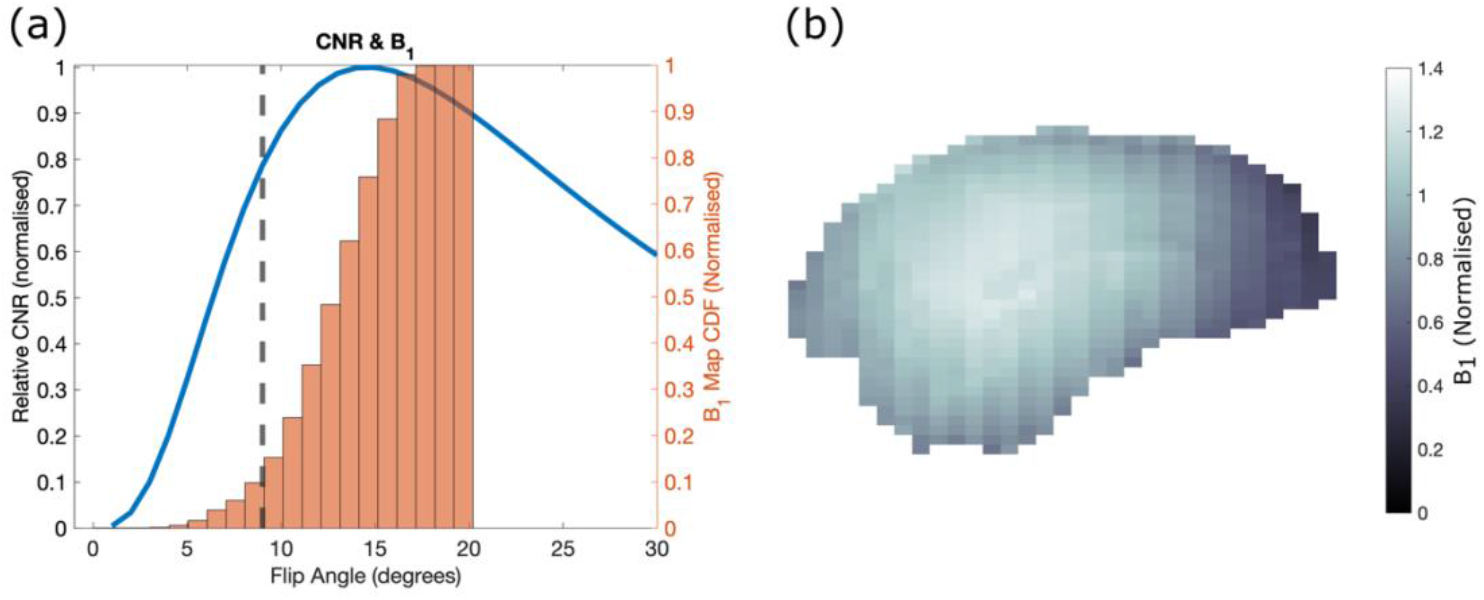
*B*_1_ inhomogeneity and relative CNR. (a) displays the theoretical DW-SSFP CNR (blue line) as a function of the flip angle for the optimised DW-SSFP sequence parameters at *b*_eff_ = 5600 s/mm^2^, alongside the cumulative distribution function (orange) of the experimental *B*_1_ map acquired in the ex vivo macaque brain. The dashed black line indicates where CNR is 80% of maximum, with under 10% of the *B*_1_ map falling below this value. (b) displays the normalised experimental *B*_1_ map (*B*_1_ inhomogeneity greatest along the anterior-posterior axis), with reduced *B*_1_ most prominent in the prefrontal cortex. As the rate of change of CNR with flip angle is slower at flip angles greater than the optimum, a small ‘overflip’ (i.e. *B*_1_ > 1) was performed in the experimental acquisition to preserve CNR.

Figure 7 displays principal diffusion direction (FA-modulated 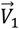), fractional anisotropy (FA) and mean diffusivity (MD) maps along all three dimensions of the macaque brain. Images demonstrate high-SNR diffusivity estimates across the entire macaque brain without geometric distortion, reflecting the quality of the acquired DW-SSFP volumes.

**Figure 7.**
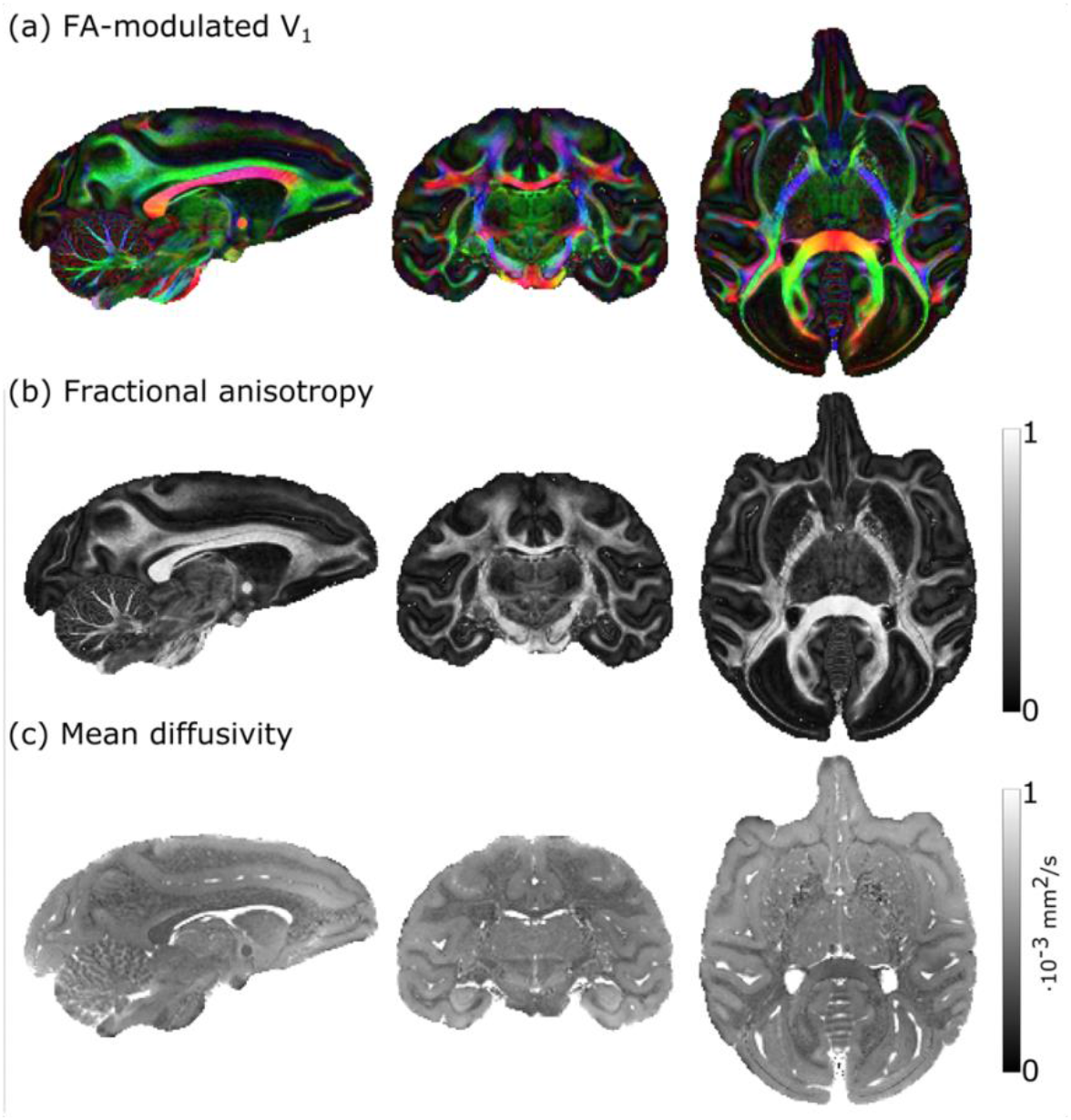
Tensor outputs. Whole brain 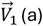 (a), fractional anisotropy (FA) (b), and mean diffusivity (MD) maps acquired in the ex vivo macaque brain. The maps display excellent spatial contrast and homogeneity across the brain sample. Images generated using a DW-SSFP signal representation incorporating a diffusion tensor as described in the methods. In (a), red = left-right, blue = superior-inferior and green = anterior-posterior. Tensor estimates based on an updated pipeline that incorporated an additional Gibbs ringing correction (Kellner et al., 2016)

Figure 8a displays whole brain fibre-orientation distribution functions, displaying excellent contrast across the brain. Figure 8b displays orientation reconstructions from the Ball and 3 Sticks outputs and CSD fODFs in the centrum semiovale (Figure 8b), a challenging crossing fibre region. Consistent 3-way crossing fibres are observed, with the CSD outputs producing fODFs indicating multiple fibre populations.

**Figure 8.**
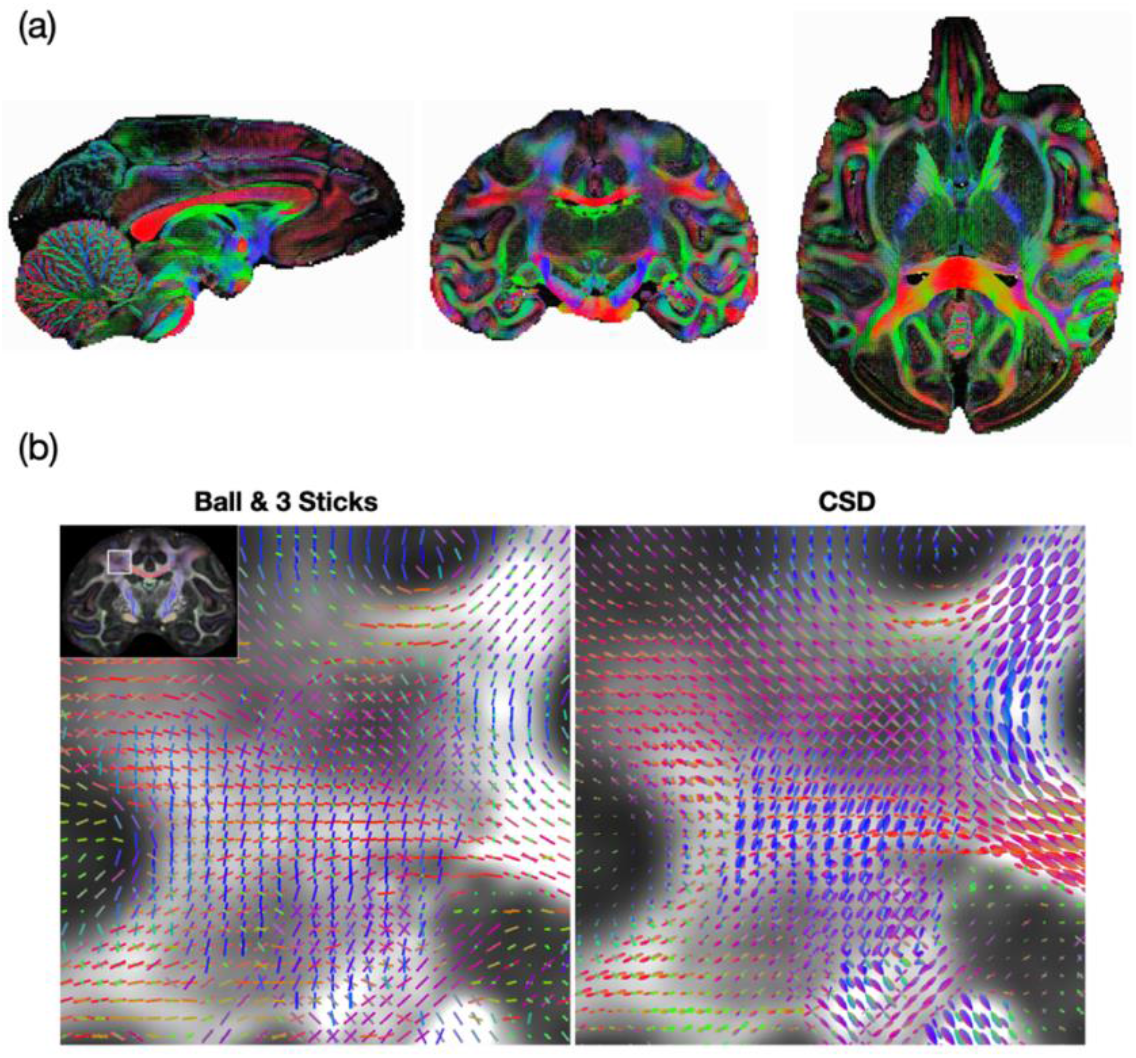
Orientation outputs. (a) displays whole-brain fibre orientation dispersion functions (fODFs) derived from the CSD investigation. (b) displays fibre orientation estimates in the centrum semiovale derived from the ball and 3 sticks (left) and CSD-SSFP (right), resolving multiple fibre populations in this challenging brain region. In (a), red = left-right, blue = superior-inferior and green = anterior-posterior.

The DW-SSFP signal has been shown to have orientation properties similar to a conventional DW-SE acquisition (McNab & Miller, 2008), making it suitable for tractography investigations. Figure 9 motivates the potential of these data to investigate brain connectivity, displaying example long range tracts reconstructed from the Ball and 3 Sticks parameter outputs.

**Figure 9.**
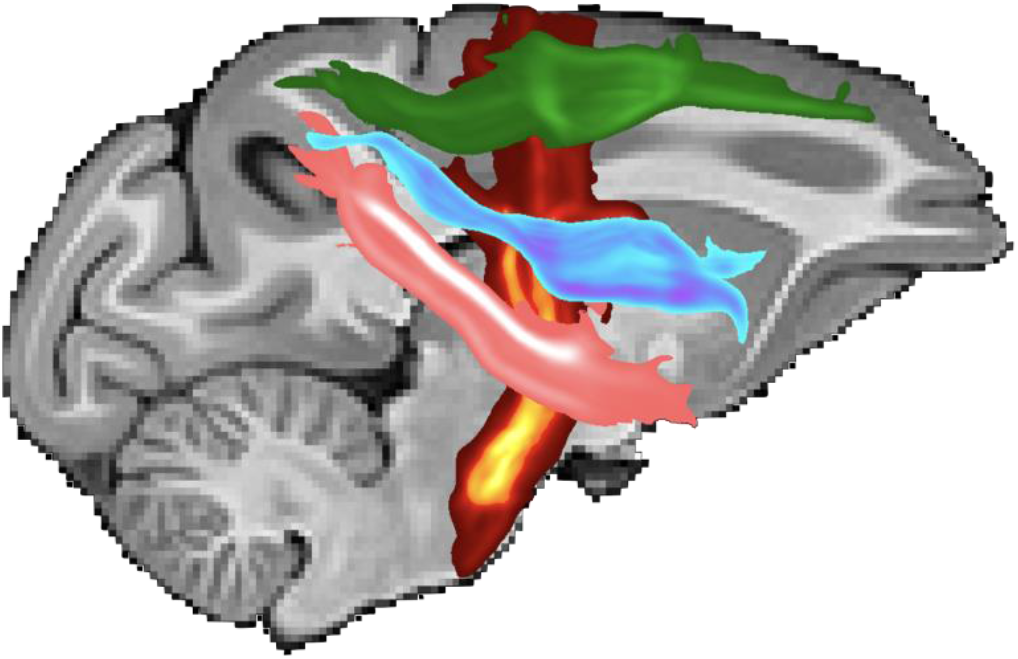
Fiber reconstructions. Sagittal maximum intensity projections of the path distributions of four bundles. Specifically, three longitudinal association tracts arising from the parietal lobe (Superior Longitudinal Fasciculus 1 - Green, Superior Longitudinal Fasciculus 3 - Blue, Middle Longitudinal Fasciculus - Pink) and a projection bundle (Corticospinal Tract - Red). Tractography performed using orientation outputs from the Ball and 3 Sticks parameter outputs using FSL-XTRACT (Assimopoulos et al., 2024; Warrington et al., 2019).

## Discussion

### Overview

The increased theoretical signal available at higher static magnetic field strengths has motivated the development of ultra-high field MRI systems to push the boundaries of achievable spatial resolution. However, additional challenges arising from changes in relaxation times, distortion sensitivity and *B*_1_ inhomogeneity limits the direct translation of diffusion MRI methods that perform well at clinical field strengths. These challenges are exacerbated in the imaging environment of fixed ex vivo tissue, requiring the development of specialised methods to overcome them.

In this work we demonstrated that the DW-SSFP sequence is one way to achieve this, with acquired diffusion imaging volumes achieving high SNR-efficiency in the challenging imaging environment of fixed ex vivo tissue at 10.5 T. Acquired data displays excellent SNR and diffusion contrast (Figures 4 and 5) suitable for connectivity investigations. A key additional benefit of DW-SSFP for high-resolution studies of precision anatomy is that it is compatible with highly-segmented (here, single-line) readouts, whereas DW-SE and DW-STE become inefficient at high levels of segmentation. As a result, our images have very low spatial distortion, similar to structural images.

### Relaxation times

We observed a substantial difference in the relaxation times from the pilot (*T*_1_ = 1350 ms and *T*_2_ = 28 ms) and measured (*T*_1_ = 1050 ms and *T*_2_ = 53 ms, and 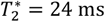) relaxation estimates in the macaque brain at the time of data acquisition. This difference likely reflects changes in relaxation times driven by the longer period of soaking in PBS for the second scan, with ∼2 months between the pilot scan and data acquisition.

A key question is whether these changes in relaxation times have a considerable impact on the achievable SNR efficiency, and how his effects the relative SNR efficiency of the different sequences. To investigate this, Figure 10 displays the findings from performing a new SNR efficiency optimisation incorporating the post-soaking relaxation times (solid lines), with optimised parameters provided in Supporting Information Table S2. Overall, the post-soaking relaxation times offer a considerable benefit for all sequences, corresponding to an average SNR efficiency increase of 274% (DW-SSFP), 523% (DW-SE) and 249% (DW-STE) across the investigated b-value range (dashed lines). Interestingly, the achievable SNR-efficiency of the DW-SSFP and DW-SE sequence are near-matched in higher b-value regimes, with a slightly higher SNR efficiency for DW-SE at low b-values (although these would offer more limited contrast given ex-vivo diffusivities). Crucially, DW-SSFP can deliver diffusion contrast whilst additionally offering the benefit of negligible image distortions, whereas DW-SE would still require segmented EPI, trading off image quality for efficiency. These findings additionally motivate sequence parameter optimisation as close as possible to data acquisition to account for any temporal variations in relaxation properties arising from the tissue fixation or soaking processes.

**Figure 10.**
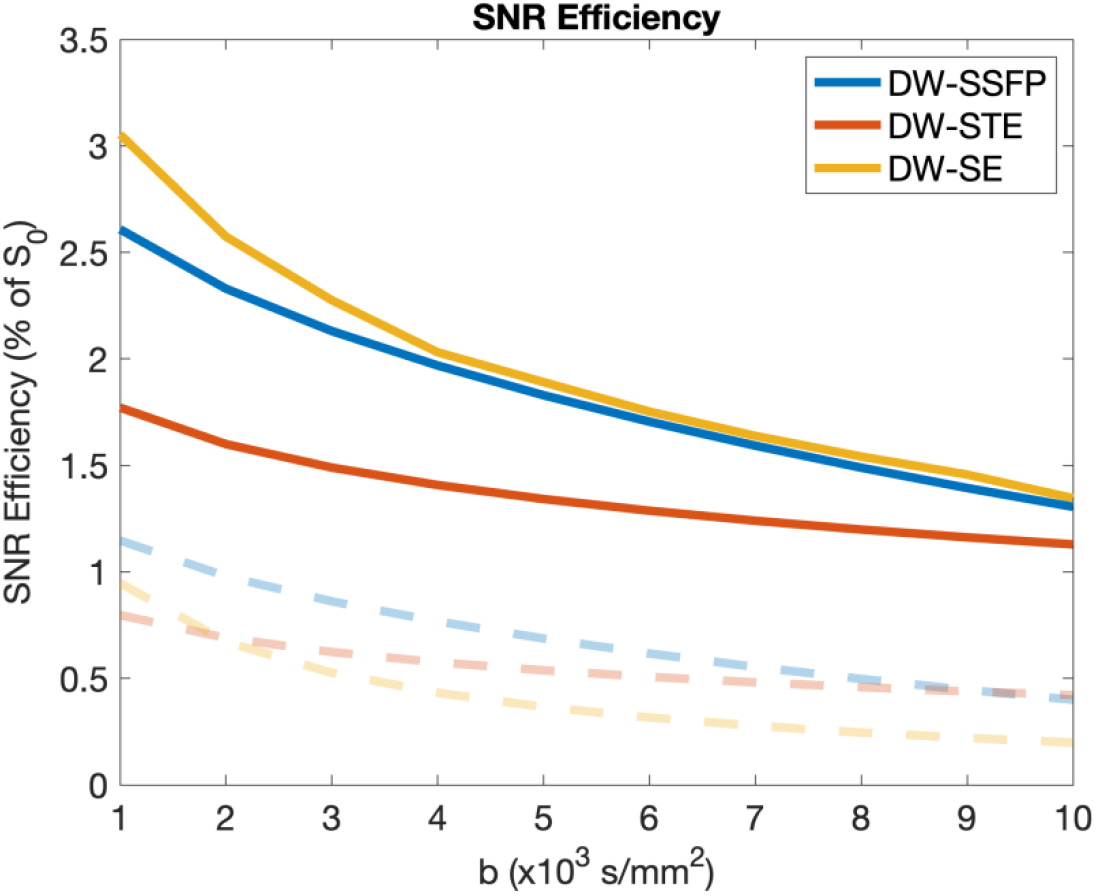
Comparison of SNR efficiency with pilot and measured relaxation times. Equivalent to the structure of Figure 3, the solid lines display the achievable SNR efficiency for the DW-SSFP, DW-SE and DW-SE sequence incorporating the relaxation times estimated at time of data acquisition (*T*_1_ = 1050 ms and *T*_2_ = 53 ms, and *T*_2_ = 24 ms). SNR efficiency estimates are higher for all three sequences, with the dashed lines displaying the equivalent SNR-efficiency estimates using the pilot relaxation values. The new relaxation values give rise to a new set of conclusions for SNR efficiency, with DW-SE performing best at low b-values (and matched to DW-SSFP at higher b-values), with DW-STE having a lower SNR efficiency.

Further simulations identified that when performing a new SNR efficiency optimisation based on the measured relaxation times (Figure 10 – solid lines) – we can achieve a further ∼17% increase in SNR efficiency relative to the parameters used for the experimental DW-SSFP acquisition. This difference indicates that there are still potential SNR-efficiency acquisition gains to be made with DW-SSFP in comparison to the proposed protocol presented here, which could translate into improved spatial resolution or increased b-values.

### Sequence optimisation - limitations

A key limitation of our SNR-efficiency optimisation framework is that it does not incorporate information about the choice of readout and overall acquisition time. For ex vivo DW-SSFP acquisitions this is typically not a problem as the sequence’s TR is short and compatible with a single line readout. This enables both a reasonable acquisition times per volume (here 13 minutes and 30 seconds) and images with negligible image distortions or blurring artefacts. DW-SE and DW-STE sequences could in theory use a single-line readout, but their long diffusion preparation times make such an acquisition incredibly inefficient and slow. For example, a single-line DW-SE protocol matching the b-value, resolution and FOV acquired here based on the SNR-optimal TR (Supporting Information Table S1) would require >18 hours per volume. DW-SE and DW-STE sequences could instead use a segmented-EPI readout, but this leads to a challenging trade-off between image distortion/blurring (favouring shorter readouts) and scan time (favouring longer readouts). This trade-off has been considered before at 3T (Miller et al., 2012), but will be considerably more acute at 10.5 T due to greater B_0_ inhomogeneity and shorter T_2_. Taken together, DW-SSFP is anticipated to provide advantages for ex vivo ultra-high field imaging even when presented with a reduced or similar SNR efficiency in comparison to the DW-SE and DW-STE sequence.

A second limitation of the proposed optimisation framework is that it does not incorporate information about the gradient duty cycle. The short TR of the DW-SSFP sequence is associated with an increased gradient duty cycle arising from the diffusion encoding module in comparison to the DW-SE and DW-STE sequence, potentially imparting limits on the maximum achievable diffusion gradient amplitude. By means of comparison, the EPI readout module of the DW-SE and DW-STE sequence imparts a greater gradient duty cycle in comparison to the DW-SSFP single line readout, impacting the available choices of readout or achievable diffusion gradient amplitude. To balance these trade-offs, we fixed the achievable *G*_max_ as equal for all investigated sequences (52 mT/m).

### Signal modelling

A frequently noted challenge of DW-SSFP is the sequence’s complicated signal-forming mechanisms (Appendix 1), with diffusion-weighting depending on local tissue relaxation times and *B*_1_ values. Here we explored several ways to model the acquired DW-SSFP data (Figures 7 and 8), identifying multiple populations in a challenging crossing-fibre region (Figure 8b), indicative of the achievable data quality at this high spatial resolution. This manuscript is, to the best of our knowledge, the first time that CSD and a Ball and 3 Sticks signal representation has been applied to DW-SSFP data.

### Future directions

The high SNR and CNR achieved with the experimental ex vivo macaque dataset (Figure 5) indicate the potential of further increasing the spatial resolution and effective b-value in future acquisitions. Figure 11 quantifies this, displaying the theoretical SNR available as a function of spatial resolution and effective b-value relative to the acquired experimental dataset. The available SNR suggests increases in spatial resolution to 0.25 mm isotropic are achievable for a similar target effective b-value, achieving SNR values > 2. For matched spatial resolution to the acquired data presented here, effective b-values above 10,000 are also achievable. Future work will explore refining the acquisition scheme to boost spatial resolution and/or increase the effective b-value.

**Figure 11.**
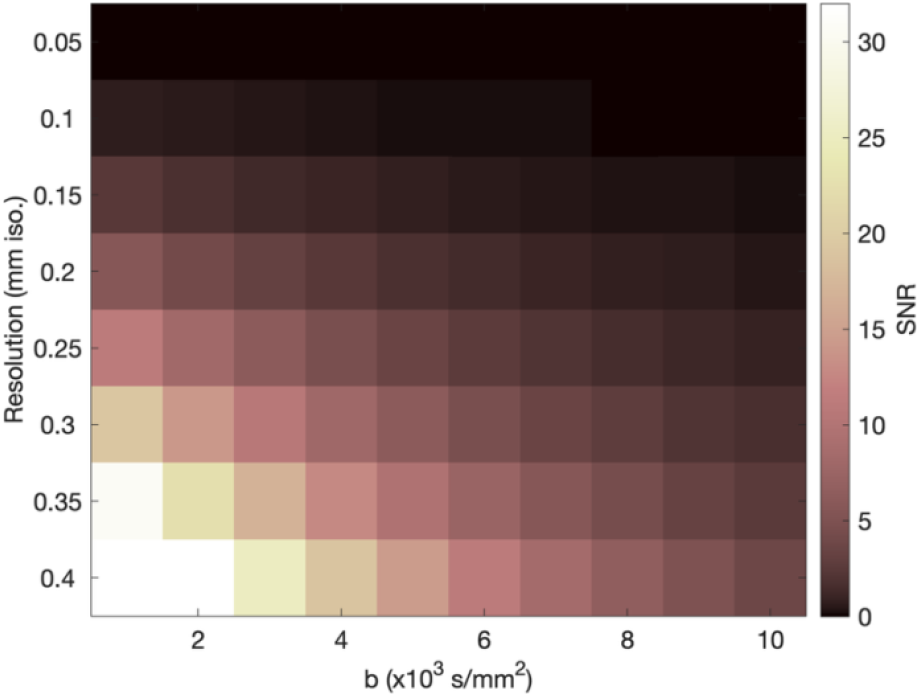
Relative DW-SSFP SNR as a function of b-value and spatial resolution. Based on the average SNR of the highest b-value shell (∼10 based on the SNR of the b_0_ volumes and average estimate of diffusion attenuation), here we display the relative SNR predicted as a function of b-value and spatial resolution. We predict SNR well above the noise level across a broad range of spatial resolutions and b-values, indicating the potential to further reduce voxel sizes and increase diffusion contrast. Here the dependency on effective b-value was modelled by identifying sequence parameters that maximise SNR efficiency (analogous to Figure 3 incorporating diffusion weighting) incorporating the experimental relaxation times, with the y-axis modelled by scaling the relative SNR by the voxel size (multiplied by (Voxel Dimension)^3^). An additional correction was performed to account for the improved SNR efficiency predicted of the optimised sequences incorporating the experimental relaxation times. No correction for 2*π* · *n* dephasing was performed.

A future aim of the Center for Mesoscale Connectomics is the acquisition of ex vivo human brain diffusion MRI data, which necessitate the use of human MRI scanners due to sample size. The findings presented in this work are anticipated to form the blueprint for the human ex vivo acquisitions, which will require further methods development to account for the impact of increased *B*_1_ inhomogeneity across the sample, and reduced control over the fixation (i.e. immersion rather than perfusion fixation with standardised brain-bank fixatives) and soaking (i.e. feasibility of soaking a whole-human brain sample) procedure.

When considering *B*_1_ inhomogeneity, work has already begun on developing a custom whole ex vivo brain imaging coil incorporating multiple transmit elements for pTx acquisitions. Due to the reduced options available for fixation and soaking, it is anticipated that imaged ex vivo human brain samples will have reduced *T*_2_ relative to the ex vivo macaque brain investigated here, requiring careful trade-offs to achieve sufficient SNR whilst keep acquisition times reasonable.

## Data and code availability

Software for the DW-SSFP, DW-SE and DW-STE SNR-efficiency optimisations, alongside scripts to recreate some of the figures in this manuscript is available at github.com/BenjaminTendler/UHF_DWSSFP. The described data will be made fully open source as part of Center for Mesoscale Connectomics (https://mesoscale-connectivity.org) and provided in a future release.

## Acknowledgements

BCT is funded by a Sir Henry Wellcome Postdoctoral Fellowship (Wellcome Trust) [222829/Z/21/Z]. KLM is funded by a Wellcome Trust Senior Research Fellowship [224573/Z/21/Z]. The work is supported by NIH grants P41EB027061 (JZ, KU, GA), UM1NS132207 (JZ, KU, CL, EY, SH), R01EB031765 (JZ) and P30DA048742 (JZ, SH).

## Contributions

**Benjamin C. Tendler:** Formal Analysis, Methodology, Software, Visualization, Writing – Original Draft Preparation. **Shaun Warrington**: Formal Analysis, Methodology, Software, Visualization. **Mohamed K. Selim**: Formal Analysis, Methodology, Software, Visualization. **Wenchuan Wu**: Methodology, Software. **Gregor Adriany**: Methodology. **Edward J. Auerbach**: Methodology. **Alexander Bratch**: Methodology, Investigation. **Hamza Farooq**: Visualization, Formal analysis. **Noam Harel**: Resources, Funding acquisition, Writing – Review & Editing. **Sarah Heilbronner**: Funding acquisition, Investigation, Project administration, Resources, Writing – Review & Editing, **Saad Jbabdi**: Funding acquisition, Methodology, Formal analysis, Software. **Steve Jungst**: Methodology. **Christophe Lenglet**: Funding acquisition, Methodology, Supervision. **Ana M.G. Manea**: Data curation, Formal analysis, Investigation, Methodology. **Steen Moeller**: Methodology, Software, Formal analysis. **Pramod K. Pisharady**: Funding acquisition, Methodology, Writing – Review & Editing. **Kamil Ugurbil**: Funding acquisition, Resources, Supervision, Project administration. **Matt Waks**: Methodology. **Essa Yacoub**: Methodology, Funding acquisition. **Stamatios N. Sotiropoulos**: Formal analysis, Methodology, Supervision, Visualization. **Karla L. Miller:** Conceptualisation, Methodology, Supervision, Writing – Review & Editing. **Jan Zimmermann:** Conceptualisation, Resources, Data curation, Investigation, Methodology, Supervision, Funding acquisition, Writing – Review & Editing.

## Appendix 1

Based on the analytical model in (Freed et al., 2001), the DW-SSFP signal associated with free Gaussian diffusion is described as:

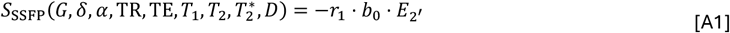

where:

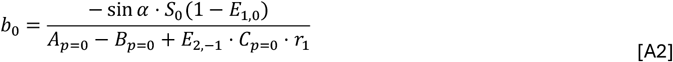

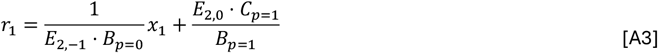

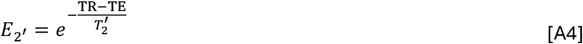

defining:

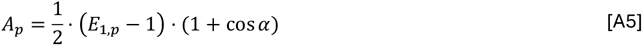

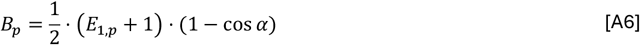

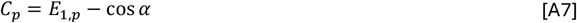

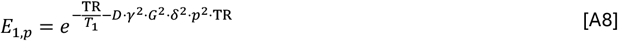

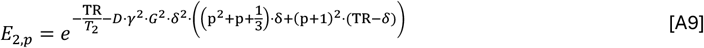

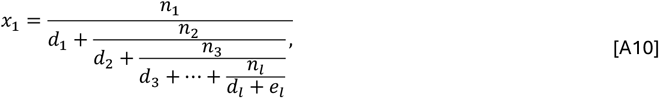

with:

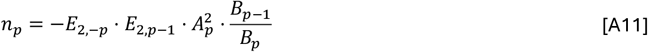

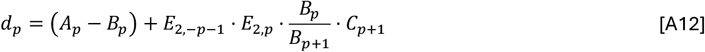

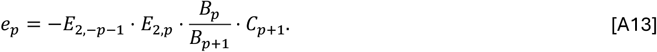

The 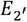 expression (Eqs. [A1] and [A4]) accounts for the impact of 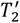 signal loss arising from the centre of the cartesian k-space readout not being aligned with the DW-SSFP echo (Figure 1a), where:

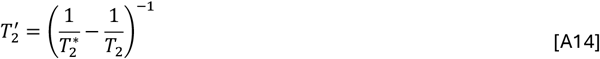

## Appendix 2

*S* for the DW-SE sequence is defined as:

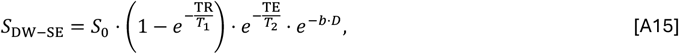

with T_acq_ equivalent to:

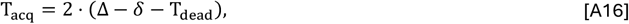

where Δ is the diffusion time.

## Appendix 3

*S* for the DW-STE sequence is defined as:

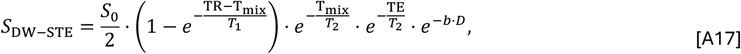

where T_mix_ is the mixing time, and T_acq_ is equivalent to:

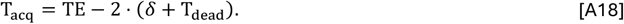

