## Supplementary Table 1 for "Diffusion-weighted steady-state free precession imaging in the ex vivo macaque brain on a 10.5T human MRI scanner"

**DW-SSFP**

| b-value (s/mm$^2$) | SNR-efficiency (% of $S_0$) | Gradient Amplitude (mT/m) | Gradient Duration (ms) | q-value (mm$^{-1}$) | Flip Angle (°) | Repetition Time (ms) | Echo Time (ms) |
|---|---|---|---|---|---|---|---|
| 1,000 s | 1.15 | 52 | 3.9 | 8.6 | 14.4 | 13.6 | 11.2 |
| 2,000 | 0.98 | 52 | 5.5 | 12.2 | 14.2 | 15.4 | 13 |
| 3,000 | 0.86 | 52 | 6.9 | 15.2 | 14.1 | 16.9 | 14.4 |
| 4,000 | 0.77 | 52 | 8.1 | 18 | 14 | 18.3 | 15.7 |
| 5,000 | 0.69 | 52 | 9.4 | 20.7 | 14.1 | 19.6 | 17 |
| 6,000 | 0.62 | 52 | 10.6 | 23.4 | 14.1 | 20.9 | 18.2 |
| 7,000 | 0.55 | 52 | 11.8 | 26.2 | 14.2 | 22.2 | 19.5 |
| 8,000 | 0.5 | 52 | 13.1 | 28.9 | 14.3 | 23.5 | 20.8 |
| 9,000 | 0.45 | 52 | 14.4 | 31.8 | 14.4 | 24.9 | 22.1 |
| 10,000 | 0.4 | 52 | 15.7 | 34.8 | 14.6 | 26.3 | 23.5 |

**DW-SE**

| b-value (s/mm$^2$) | SNR-efficiency (% of $S_0$) | Gradient Amplitude (mT/m) | Gradient Duration (ms) | Diffusion Time (ms) | Echo time (ms) | Readout Duration (ms) | Repetition Time (ms) |
|---|---|---|---|---|---|---|---|
| 1,000 | 0.95 | 52 | 14.6 | 29 | 58 | 18.7 | 1700 |
| 2,000 | 0.67 | 52 | 19.3 | 34.1 | 68.2 | 19.5 | 1700 |
| 3,000 | 0.53 | 52 | 22.7 | 37.7 | 75.4 | 20 | 1700 |
| 4,000 | 0.43 | 52 | 25.4 | 40.6 | 81.1 | 20.4 | 1700 |
| 5,000 | 0.37 | 52 | 27.7 | 43 | 86 | 20.6 | 1700 |
| 6,000 | 0.32 | 52 | 29.7 | 45.1 | 90.2 | 20.9 | 1700 |
| 7,000 | 0.28 | 52 | 31.5 | 47 | 94 | 21 | 1700 |
| 8,000 | 0.25 | 52 | 33.1 | 48.7 | 97.4 | 21.2 | 1700 |
| 9,000 | 0.22 | 52 | 34.6 | 50.3 | 101 | 21.3 | 1700 |
| 10,000 | 0.2 | 52 | 36 | 51.8 | 104 | 21.5 | 1700 |

**DW-STE**

| b-value (s/mm$^2$) | SNR-efficiency (% of $S_0$) | Gradient Amplitude (mT/m) | Gradient Duration (ms) | Mixing Time (ms) | Echo time (ms) | Readout Duration (ms) | Repetition Time (ms) |
|---|---|---|---|---|---|---|---|
| 1,000 | 0.8 | 52 | 5.2 | 174 | 34.7 | 14.3 | 2060 |
| 2,000 | 0.69 | 52 | 6.6 | 223 | 37.3 | 14.2 | 2160 |
| 3,000 | 0.62 | 52 | 7.5 | 258 | 39.2 | 14.2 | 2230 |
| 4,000 | 0.58 | 52 | 8.2 | 286 | 40.7 | 14.2 | 2280 |
| 5,000 | 0.54 | 52 | 8.9 | 310 | 41.9 | 14.2 | 2320 |
| 6,000 | 0.51 | 52 | 9.4 | 331 | 43 | 14.2 | 2360 |
| 7,000 | 0.48 | 52 | 9.9 | 350 | 44 | 14.2 | 2390 |
| 8,000 | 0.46 | 52 | 10.3 | 368 | 44.9 | 14.2 | 2430 |
| 9,000 | 0.44 | 52 | 10.7 | 383 | 45.7 | 14.2 | 2450 |
| 10,000 | 0.42 | 52 | 11.1 | 398 | 46.4 | 14.2 | 2480 |

Table S1: **Estimated parameters from the SNR-efficiency optimisations.** Here we provide the estimated sequence parameters from the SNR-efficiency optimisations for the DW-SSFP sequence (top), DW-SE sequence (middle), and DW-STE sequence (bottom). Note that no correction for $2\pi \cdot n$ dephasing has been performed for the DW-SSFP sequence.

**DW-SSFP**

| b-value (s/mm$^2$) | SNR-efficiency (% of $S_0$) | Gradient Amplitude (mT/m) | Gradient Duration (ms) | q-value (mm$^{-1}$) | Flip Angle (°) | Repetition Time (ms) | Echo Time (ms) |
|---|---|---|---|---|---|---|---|
| 1,000 s | 2.61 | 52 | 4.6 | 10.2 | 21.4 | 17.1 | 13.3 |
| 2,000 | 2.33 | 52 | 6.5 | 14.4 | 21.2 | 19.5 | 15.5 |
| 3,000 | 2.13 | 52 | 8.1 | 18 | 21.2 | 21.4 | 17.2 |
| 4,000 | 1.97 | 52 | 9.6 | 21.2 | 21.2 | 23.1 | 18.8 |
| 5,000 | 1.83 | 52 | 11 | 24.3 | 21.2 | 24.7 | 20.3 |
| 6,000 | 1.71 | 52 | 12.4 | 27.4 | 21.3 | 26.3 | 21.8 |
| 7,000 | 1.59 | 52 | 13.7 | 30.4 | 21.5 | 27.8 | 23.3 |
| 8,000 | 1.49 | 52 | 15.1 | 33.5 | 21.6 | 29.4 | 24.8 |
| 9,000 | 1.39 | 52 | 16.5 | 36.6 | 21.8 | 31 | 26.3 |
| 10,000 | 1.31 | 52 | 17.9 | 39.7 | 22 | 32.6 | 27.8 |

**DW-SE**

| b-value (s/mm$^2$) | SNR-efficiency (% of $S_0$) | Gradient Amplitude (mT/m) | Gradient Duration (ms) | Diffusion Time (ms) | Echo time (ms) | Readout Duration (ms) | Repetition Time (ms) |
|---|---|---|---|---|---|---|---|
| 1,000 | 3.06 | 51.9 | 13.4 | 33.4 | 66.8 | 30 | 1040 |
| 2,000 | 2.58 | 51.8 | 18 | 38 | 76 | 30 | 1080 |
| 3,000 | 2.29 | 52 | 21.3 | 41.3 | 82.6 | 30 | 1100 |
| 4,000 | 2.07 | 52 | 24 | 44 | 87.9 | 30 | 1110 |
| 5,000 | 1.9 | 52 | 26.2 | 46.2 | 92.5 | 30 | 1130 |
| 6,000 | 1.77 | 52 | 28.3 | 48.3 | 96.5 | 30 | 1140 |
| 7,000 | 1.65 | 52 | 30.1 | 50.1 | 100 | 30 | 1150 |
| 8,000 | 1.55 | 52 | 31.7 | 51.7 | 103 | 30 | 1160 |
| 9,000 | 1.47 | 52 | 33.2 | 53.2 | 106 | 30 | 1160 |
| 10,000 | 1.39 | 52 | 34.6 | 54.6 | 109 | 30 | 1170 |

**DW-STE**

| b-value (s/mm$^2$) | SNR-efficiency (% of $S_0$) | Gradient Amplitude (mT/m) | Gradient Duration (ms) | Mixing Time (ms) | Echo time (ms) | Readout Duration (ms) | Repetition Time (ms) |
|---|---|---|---|---|---|---|---|
| 1,000 | 1.77 | 52 | 7.1 | 77.6 | 52 | 27.7 | 1480 |
| 2,000 | 1.6 | 52 | 9 | 103 | 55.6 | 27.7 | 1540 |
| 3,000 | 1.49 | 52 | 10.3 | 122 | 58.2 | 27.6 | 1570 |
| 4,000 | 1.41 | 52 | 11.3 | 136 | 60.2 | 27.6 | 1600 |
| 5,000 | 1.34 | 52 | 12.1 | 149 | 61.9 | 27.6 | 1630 |
| 6,000 | 1.29 | 52 | 12.9 | 160 | 63.4 | 27.6 | 1650 |
| 7,000 | 1.24 | 52 | 13.5 | 170 | 64.7 | 27.6 | 1670 |
| 8,000 | 1.2 | 52 | 14.1 | 178 | 65.9 | 27.6 | 1680 |
| 9,000 | 1.16 | 52 | 14.7 | 187 | 67 | 27.6 | 1700 |
| 10,000 | 1.13 | 52 | 15.2 | 194 | 68 | 27.6 | 1720 |

Table S2: **Estimated parameters from the SNR-efficiency optimisations with updated relaxation times.** Here we provide the estimated sequence parameters from the SNR-efficiency optimisations for the DW-SSFP sequence (top), DW-SE sequence (middle), and DW-STE sequence (bottom). Here ($T_1 = 1050$ ms and $T_2 = 53$ ms, and $T_2^* = 24$ ms). Note that no correction for $2\pi \cdot n$ dephasing has been performed for the DW-SSFP sequence.
